# Pathogen and pest communities in agroecosystems across climate gradients: Anticipating future challenges in the highland tropics

**DOI:** 10.1101/2025.01.08.631994

**Authors:** Romaric A. Mouafo-Tchinda, Aaron I. Plex Sulá, Berea A. Etherton, Joshua S. Okonya, Gloria Valentine Nakato, Yanru Xing, Jacobo Robledo, Ashish Adhikari, Guy Blomme, Déo Kantungeko, Anastase Nduwayezu, Jan F. Kreuze, Jürgen Kroschel, James P. Legg, Karen A. Garrett

## Abstract

**CONTEXT:** Tropical cropping systems must adapt to the current and future geographic distribution of pathogen and pest communities. An important research gap is how climate change may shift the distribution of pathogens and pests in tropical lowlands and highlands.

**OBJECTIVE:** We evaluated the current geographic risk of 27 pathogens and pests in the production of four food security crops (banana, cassava, potato, and sweetpotato) in the Great Lakes region of Africa, and the potential future risk under climate change. Models for each pathogen and pest indicate the potential for changes in geographic distribution, with model fit indicating the potential for decision support systems to facilitate management.

**METHODS:** First, cropland connectivity analysis identified locations likely important in the spread of crop-specific pathogens and pests, such as locations in Rwanda and Burundi. Second, we surveyed the 27 economically important pathogens and pests in Rwanda and Burundi, mapping the distribution of each across climate gradients and quantifying patterns of association. Third, we used machine learning to develop models of each species as a function of environmental variables, including host landscape variables. We also evaluated the increase in temperature across altitudes under future climate change scenarios in this region.

**RESULTS AND CONCLUSIONS:** Among the ten machine-learning algorithms evaluated, random forests and support vector machines generally performed best for predicting severity and infestation. Host landscape variables were useful predictors for some species. Based on climate matching, 44% of the pathogens and pests could become more common with warmer temperatures at higher altitudes, while 17% may become less common.

**SIGNIFICANCE:** This study indicates adaptation priorities for crop health in a region with multiple challenges to agricultural sustainability. The models developed here also indicate which species may have more potential and relevance for future development of pathogen and pest forecasts.

## 1. Introduction

### 1.1. Impact of climate and crop landscape structure on plant pathogens and pests

Agroecosystems in the tropics, such as the Great Lakes region of Africa, are often challenged by climate variability and societal dynamics including demographic pressures, socio-political instability, and resource-related conflict (Ogot and Niane, 1984; Ong’ayo, 2018). Climate change is likely to affect agricultural productivity and exacerbate the proliferation of some pathogens and pests in this region (Ebregt *et al*., 2005; Ferdu and Solbreck, 2010; Okonya and Kroschel, 2013; Gallego-Tévar *et al*., 2024). Ecosystems can be reshaped by new temperature and precipitation patterns, driving pathogens and pests to new locations (Sparks *et al*., 2014; Cilas *et al*., 2016; Garrett *et al*., 2022a; Garrett *et al*., 2022b). The interactions among climate heterogeneity, the geographic structure of agroecosystems, and pathogens and pests in tropical regions such as the Great Lakes region represent a major knowledge gap. Research has examined how climate affects the dynamics of specific pathogens and pests (Kroschel *et al*., 2014; Adhikari *et al*., 2015; Blomme *et al*., 2020). However, field studies that allow for simultaneous comparisons of many different pathogens and pests along climate gradients are rare, and little is known about how regional factors, such as the geographic structure of agroecoystems in the Great Lakes region, exacerbate risks. It is an open question how pathogen and pest communities are distributed across altitudes in this region, what variables are predictive of pathogen and pest community structure, and how their distribution may shift under climate change.

Climate matching studies identify which locations have similar climate conditions, such that they may be equally suitable for a species; ecological communities may move following changes in where suitable climate conditions are found in the future (Colwell *et al*., 2008; Singh *et al*., 2023; Chevalier *et al*., 2024). For example, pathogen and pest communities now at hotter, lower altitudes may move to what are currently cooler, higher altitudes if there is warming due to climate change. The dynamics of pathogens and pests in response to climate have been widely studied (Hodkinson, 2005; Pautasso *et al*., 2012; Juroszek *et al*., 2022; Mishra *et al*., 2024). Higher altitudes are commonly associated with lower species richness, disease severity, and pest damage (Hodkinson, 2005; Blomme *et al*., 2020). Since altitude is typically negatively correlated with temperature (Poveda *et al*., 2012), altitude can be a useful proxy when evaluating the effect of temperature on plant disease and pest damage. For example, previous studies in South America and East Africa predicted that rising temperatures due to climate change will modify the geographic distribution of pathogens and pests, increasing plant disease severity or pest damage at higher altitudes (Poveda *et al*., 2012; Blomme *et al*., 2020). New climate conditions favorable to certain pathogens and pests are likely to occur at higher altitudes, disrupting ecological balances and exposing higher-altitude plants and ecosystems to new risks.

The geographic distribution of crop production can also influence geographic patterns of disease severity and pest damage. Crop landscape structure has been well documented to influence crop damage from diseases and pests (Thies and Tscharntke, 1999; Poveda *et al*., 2012). Analysis of the potential invasion network for pathogens and pests in crop landscapes, such as cropland connectivity analysis, has been used to identify candidate priority locations for surveillance and management of host-specific pathogens and pests (Xing *et al*., 2020; Andersen Onofre *et al*., 2021; Buddenhagen *et al*., 2022). An analysis of cropland geographic structure indicates the geographic distribution of potential spread of pathogens and pests. Higher host plant availability generally increases the local risk of host-specific pathogen and pest establishment, while greater host connectivity can also increase the risk of spread and establishment.

### 1.2. Disaster plant pathology: Vulnerabilities and solutions in the Great Lakes region

Many families living in the Great Lakes region rely on food security crops such as banana and plantain, cassava, potato, and sweetpotato. However, many households remain food insecure because of declining agricultural productivity, driven by frequent disasters in the region, including conflicts, health emergencies, and natural disasters that displace populations and disrupt agriculture (Ochieng *et al*., 2014; Ochieng *et al*., 2015; Ong’ayo, 2018). These challenges are compounded by the cascading effects of high prevalence of plant pests and diseases, low agricultural input use, and loss of harvest and assets (Kroschel *et al*., 2014; McEwan, 2016; Almekinders *et al*., 2019). The Rwandese genocide in 1994, the civil war in Burundi from 1993 to 2005 and also in 2015, and the persistent conflicts in Eastern DR Congo are particularly important humanitarian disasters in the Great Lakes region, which have left an indelible mark (Ong’ayo, 2018). Health emergencies and natural disasters, such as severe droughts in Ethiopia, Kenya, and Uganda (2015-2016) and devastating floods in Rwanda, Uganda, and DR Congo (2020), have severely restricted agricultural practices such as effective surveillance and management of fields, causing considerable damage to infrastructure and cropland (Ochieng *et al*., 2014; Ochieng *et al*., 2015).

Disaster plant pathology, integrating concepts from plant pathology and disaster science, addresses the risks and potential impact of plant diseases before, during, and after disaster events that threaten agricultural systems and ecosystems (Etherton *et al*., 2024; Mouafo-Tchinda *et al*., 2024). It focuses on how disturbances modify host-pathogen dynamics and increase damage and the spread of pathogens, and on the development of strategies for better managing plant health. A combination of global and national research efforts has been adopted to manage the impacts that these disasters, pathogens, and pests may have on agroecological systems (RTB, 2013; Science Group Project on Accelerated Varietal Improvement and Seed Systems in Africa, 2024). For example, disease control programs which encourage farmers to recognize and destroy diseased plants, and disease-resistant crop breeding programs are currently underway across the Great Lakes region.

When models of pathogen and pest responses to environmental variables are available, decision-support tools can be developed to predict where specific disease- and pest-resistant varieties will be needed in the long-run, and in the short-run whether likely economic injury levels motivate management actions by growers (Kaundal *et al*., 2006; Carisse and Fall, 2021; Fenu and Malloci, 2021). Pest and disease forecasting can reduce production costs, pesticide use, and environmental pollution, protecting the health of farmers and consumers. Machine learning can be used to analyze high volume datasets, and improve the predictive power, accuracy, and efficiency for diseases and pests (Kaundal et al., 2006; Garrett et al., 2022a). Implementing decision support systems in low-resource countries such as those in the Great Lakes region is challenging; in addition to the often-limited availability of high-resolution local weather data as input to models, field observations and georeferenced datasets for the many important pathogen and pest species are often unavailable for constructing predictive models.

### 1.3. Objectives

In this study, we evaluated the geographic distribution and risk factors for 27 pathogens and pests in the production of four food security crops — banana (including plantain), cassava, potato, and sweetpotato — in the Great Lakes region of Africa, including the potential future risk under climate change. The first objective was to analyze the geographic risk of host-specific pathogen and pest spread based on host landscape connectivity for these crops. The second objective was to characterize the seasonal and geographic occurrence and associations of 27 economically important pathogens and pests affecting these crops, through field surveys conducted in two key locations in the region, Rwanda and Burundi. Part of this dataset has been used to characterize the prevalence of potato viruses and vectors (Okonya *et al*., 2021) and the occurrence of banana pests and diseases across altitudes (Nakato *et al*., 2023). The third objective was to predict pathogen and pest severity/infestation based on climate variables and host landscape structure using machine learning, and to assess which pathogens and pests are likely to experience altitudinal range shifts under projected changes in temperature. The field surveys were implemented along environmental gradients to compare lower- and higher-altitude pathogen and pest communities, where lower-altitude communities have the potential to move toward higher altitudes in the future. The simultaneous study of these 27 pathogens and pests allows a unique direct comparison of their associations with environmental predictors. These analyses can also support decision-making by (i) national agricultural agencies that plan geographic surveillance strategies for economically important pathogens and pests, (ii) local agricultural research communities and international partners as they prioritize efforts to improve current and future mitigation responses to pathogens and pests, and (iii) farmer communities that need to be prepared for changes in pathogen and pest dynamics due to natural or human-driven disasters, including changing climates.

## 2. Materials and methods

### 2.1. Landscape structures influencing pathogen and pest risk

#### 2.1.1. Identifying locations with high cropland connectivity

The spatial pattern of host availability is a key ecological trait that influences the spread of host-specific pathogens and pests. We analyzed the connectivity of host landscapes in the Great Lakes region to map the potential establishment and regional spread of host-specific pathogens and pests, to identify candidate priority locations for spatially targeted surveillance in four food security crops — banana (including plantain), cassava, potato, and sweetpotato. This cropland connectivity analysis assesses how crop production is connected in a landscape, based on host availability and physical proximity, in a gravity model (Jongejans *et al*., 2015). Cropland connectivity analysis identifies potential habitat networks through which pests and pathogens can spread, a first approximation to epidemic or invasion networks, which can be refined as more data become available (Margosian *et al*., 2009; Xing *et al*., 2020; Andersen Onofre *et al*., 2021).

For each crop, we used publicly available estimates of the cropland harvested area (or cropland density) across sub-Saharan Africa in 2017 (International Food Policy Research Institute, 2020) as a proxy for host availability to host-specific pathogens and pests. Maps of cropland density, aggregated at a spatial resolution of 10-min (or grid cells of image 9.27 km × 9.27 km), were used to evaluate the potential epidemic or invasion network using a gravity model (Xing *et al*., 2020). Here the gravity model incorporated two pathogen or pest dispersal kernels: 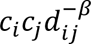 for the inverse power law model, and 𝑐_𝑖_𝑐_𝑗_𝑒^−^*^𝛾𝑑𝑖𝑗^* for the negative exponential model. In each dispersal model, 𝑑_𝑖𝑗_ is the Vincenty ellipsoid distance between a pair of cropland nodes (locations) 𝑖 and 𝑗, 𝑐_𝑖_ is the cropland density for node 𝑖, and 𝛽 and 𝛾 are dispersal parameters describing the relative likelihood of pathogen or pest movement across 𝑑_𝑖𝑗_(Xing *et al*., 2020; Andersen Onofre *et al*., 2021).

We calculated a cropland connectivity risk index (CCRI) (Xing *et al*., 2020) for each location with crop production in the region as a measure of its likely importance in epidemic or invasion networks. The CCRI was the weighted mean of four network metrics (node centralities), evaluated using the geohabnet package v2.0.0 (Keshav *et al*., 2024) in R v4.4.0 (R Core Team, 2024), averaged across the results from a sensitivity analyses for a range of dispersal parameters, as described by Mouafo-Tchinda *et al*. (2024). Two CCRI formulations were used to capture key facets of node connectivity. The first CCRI considered (“CCRI-betweenness”) was the mean of betweenness centrality weighted by 1/2, node strength weighted by 1/6, eigenvector centrality weighted by 1/6, and the sum of nearest neighbors’ degrees weighted by 1/6. The second, “CCRI-closeness”, considered 1/2 closeness centrality instead of betweenness centrality, with the same weights for the other metrics. We emphasized betweenness or closeness centrality to account for, respectively, the importance of a node as a bridge between parts of the network and the relative facility with which a location can be reached from all other locations in the network.

Habitat connectivity analysis using the geohabnet package (Keshav *et al*., 2024) is part of the R2M toolbox for rapid risk assessment supporting mitigation of pathogens and pests (<u>garrettlab.com/r2m</u>). Other examples of applications of habitat connectivity and R2M tools, in addition to those discussed elsewhere in this paper, illustrate how these tools can be used in an integrated analysis (Andersen Onofre *et al*., 2019; Garrett, 2021; Etherton *et al*., 2023; Mouafo-Tchinda *et al*., 2024; Etherton *et al*., 2025).

#### 2.1.2. Associations among altitude, crop landscape structure, and climate

We characterized associations among altitude, crop landscape structure, and climate in the Great Lakes region. The map of altitudes was retrieved from the Shuttle Radar Topography Mission (SRTM) database, using the geodata package v0.5-8 (Hijmans *et al*., 2023) in R. We also retrieved the 19 bioclimatic variables for the region from Worldclim database v2.1 (Fick and Hijmans, 2017) using the geodata package version 0.5-8 (Hijmans *et al*., 2023) and the terra package v1.8-0 (Hijmans, 2024) in R. For crop landscape structure variables, we used the raster maps of cropland density and connectivity (CCRI-closeness) generated in section 2.1.1, using the terra package (Hijmans *et al*., 2022). We assessed the associations between altitude, bioclimatic variables, and cropland structure by calculating the Pearson correlations between these variables using the metan package v1.18.0 (Olivoto and Lúcio, 2020) in R.

### 2.2. Pathogen and pest communities

#### 2.2.1. Field sampling and spatiotemporal structure of pathogen and pest communities

For each crop species, we identified fields for sampling along four survey transects in Rwanda and Burundi, each transect representing an altitude gradient. In Burundi, the field surveys included cropland around Lake Tanganyika in the Rusizi watershed, representing the Bubanza, Bujumbura Rural, Cibitoke, and Muramvya provinces (Fig. 1C). In Rwanda, the field surveys were in major cropland around Lake Kivu in the Ruhengeri watershed, which includes part of the Northern and Western provinces (Fig. 1C). The survey locations represented major crop landscapes in both countries, encompassing a mixture of cropping systems and a range in altitude from 900 to 2800 m.a.s.l. (Fig. 1a-b) (Kroschel *et al*., 2014). Fields for sampling were selected at regular intervals along the transects regardless of whether diseases or pest damage were apparent. The number of fields sampled in Burundi and Rwanda was 189 and 103 for banana, 78 and 116 for potato, and 107 and 104 for sweetpotato, respectively. For cassava, sampling was only carried out in 50 fields in Burundi, with the second Burundi survey not conducted due to civil unrest, and Rwanda was not sampled because few farmers grew cassava in the Rwanda study area.

**Fig. 1.**
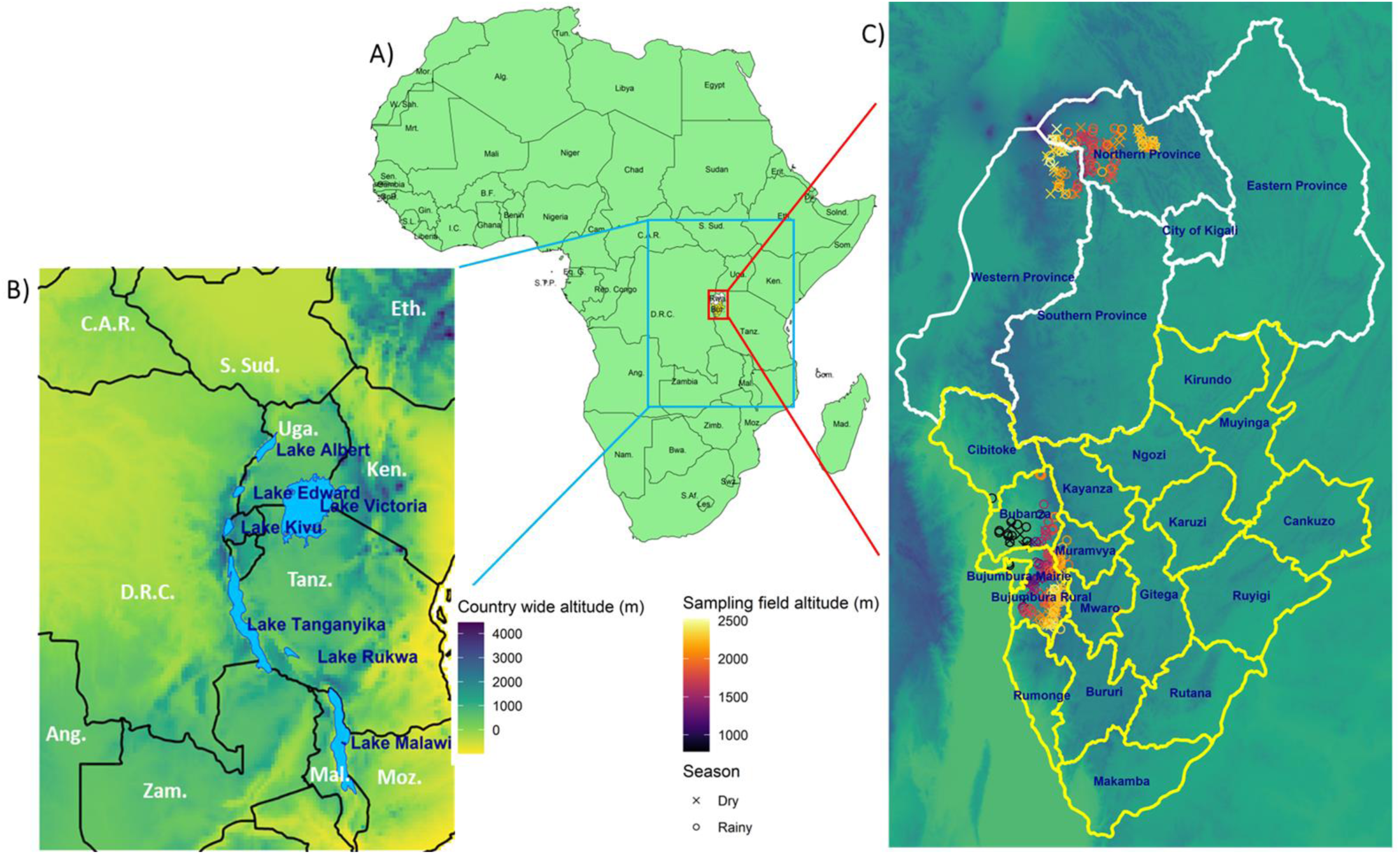
Location of fields of banana, cassava, potato, and sweetpotato where pathogen and pest surveys were carried out in Africa (A), in the Great Lakes region (B), in transects across altitude gradients, in Rwanda and Burundi (C). White lines in (C) indicate the province boundaries of Rwanda and yellow lines indicate the province boundaries of Burundi. Crosses and circles represent locations surveyed during the dry and rainy season, respectively. The complete surveillance effort included 412 crop fields.

Sampling was conducted in each watershed during two cropping seasons, dry and rainy. In both countries, two rainy seasons occur: one from March to May and the other from September to December (WorldData.info, 2024). The dry seasons occur from June to August and January to February (WorldData.info, 2024). For each field sampled, we evaluated crop-specific pest or disease intensity (Table 2) based on prevalence in the region, disease severity, infestation rate and intensity, and/or the number of pests, using methods as described by Nakato *et al*. (2023), Okonya *et al*. (2021), and Legg *et al*. (2009). A total of 27 economically important pathogens and pests in the region were included in this study, including 6 for banana, 6 for cassava, 7 for potato and 8 for sweetpotato (Table 2). Here prevalence refers to the presence or absence of a pest or disease in a field, while disease severity, pest infestation intensity or infestation rate is the percentage of leaves or plants damaged by a disease or pest. For infestation rate, infestation intensity, and severity, twenty plants per field, selected in a systematic pattern, were assessed for visual symptoms of damage by pests and diseases. The percentage of plants infested was noted as the infestation rate. The percentage of symptomatic leaves per plant was noted as infestation intensity for pests and severity for diseases. For damage by some virus diseases and pests such as cassava mosaic disease (CMD) and cassava green mite (CGM), crop damage was scored from 1 (representing no symptoms) to 5 (the most severe) (Sseruwagi *et al*., 2004; Legg *et al*., 2009; Okonya *et al*., 2021). The disease or pest damage score measures the severity or infestation intensity in a field, based on visual assessments and a predefined scale. Thirty plants were randomly sampled at regular intervals along the season and damage was assessed by scoring each plant on a scale of 1 to 5 as described by Legg *et al*. (2009).

**Table 1.** Support vector machine (SVM) and random forest (RF) performance for predicting pathogen and pest levels in smallholder farms in Burundi and Rwanda, for selected species with higher model performance. The response variable was severity for the pathogens and infestation intensity for pests, except for infestation rate for sweetpotato aphids.

|  |  |  | SVM |  |  |  | Random Forest |  |  |  |  |  |
| --- | --- | --- | --- | --- | --- | --- | --- | --- | --- | --- | --- | --- |
|  |  |  | Training |  |  | Prediction |  | Training |  |  | Prediction |  |
| Crop | Pathogen/pest | Timeframe | %RMSE | R-squared | %MAE | %RMSE | R-squared | %RMSE | R-squared | %MAE | %RMSE | R-squared |
| Banana | *P. nigronevosa* | Monthly | 18 | 0.63 | 13 | 21 | 0.56 | 16 | 0.72 | 12 | 18 | 0.66 |
|  |  | Quarterly | 18 | 0.63 | 13 | 21 | 0.56 | 16 | 0.72 | 12 | 19 | 0.64 |
|  |  | Annual | 19 | 0.63 | 14 | 21 | 0.55 | 16 | 0.71 | 11 | 19 | 0.63 |
|  | Banana bunchy top virus | Monthly | 7 | 0.22 | 3 | 4 | 0.10 | 8 | 0.25 | 4 | 5 | 0.12 |
|  |  | Quarterly | 7 | 0.22 | 3 | 4 | 0.11 | 8 | 0.23 | 4 | 5 | 0.13 |
|  |  | Annual | 7 | 0.20 | 3 | 4 | 0.09 | 8 | 0.24 | 4 | 5 | 0.13 |
|  | Plant parasitic nematodes | Monthly | 11 | 0.42 | 6 | 15 | 0.29 | 10 | 0.42 | 6 | 14 | 0.31 |
|  |  | Quarterly | 11 | 0.40 | 6 | 15 | 0.25 | 11 | 0.37 | 7 | 14 | 0.32 |
|  |  | Annual | 11 | 0.39 | 6 | 15 | 0.23 | 11 | 0.37 | 7 | 14 | 0.29 |
| Potato | Leaf miner fly | Monthly | 18 | 0.69 | 13 | 16 | 0.71 | 18 | 0.70 | 12 | 16 | 0.71 |
|  |  | Quarterly | 18 | 0.69 | 13 | 16 | 0.71 | 18 | 0.71 | 11 | 16 | 0.69 |
|  |  | Annual | 18 | 0.70 | 13 | 16 | 0.72 | 17 | 0.72 | 11 | 17 | 0.69 |
|  | *Ralstonia solanacearum* | Monthly | 14 | 0.30 | 10 | 14 | 0.13 | 13 | 0.40 | 10 | 15 | 0.12 |
|  |  | Quarterly | 15 | 0.28 | 10 | 14 | 0.13 | 13 | 0.41 | 10 | 15 | 0.12 |
|  |  | Annual | 15 | 0.28 | 11 | 14 | 0.16 | 12 | 0.39 | 10 | 15 | 0.13 |
|  | Potato viruses | Monthly | 20 | 0.31 | 16 | 21 | 0.34 | 21 | 0.28 | 17 | 21 | 0.38 |
|  |  | Quarterly | 20 | 0.32 | 16 | 21 | 0.33 | 20 | 0.31 | 16 | 21 | 0.38 |
|  |  | Annual | 20 | 0.31 | 16 | 21 | 0.34 | 21 | 0.30 | 17 | 21 | 0.37 |
| Sweet potato | *Alternaria bataticola* | Monthly | 15 | 0.35 | 9 | 10 | 0.42 | 14 | 0.39 | 9 | 10 | 0.39 |
|  |  | Quarterly | 14 | 0.41 | 9 | 9 | 0.43 | 13 | 0.45 | 8 | 9 | 0.42 |
|  |  | Annual | 15 | 0.35 | 8 | 9 | 0.47 | 14 | 0.38 | 9 | 9 | 0.43 |
|  | Sweetpotato weevils (*Cylas* spp.) | Monthly | 19 | 0.44 | 10 | 25 | 0.26 | 18 | 0.38 | 10 | 22 | 0.44 |
|  |  | Quarterly | 19 | 0.42 | 10 | 26 | 0.21 | 19 | 0.38 | 11 | 22 | 0.41 |
|  |  | Annual | 20 | 0.41 | 10 | 25 | 0.27 | 19 | 0.36 | 11 | 23 | 0.40 |
|  | Sweetpotato aphids | Monthly | 14 | 0.60 | 6 | 9 | 0.60 | 10 | 0.53 | 5 | 7 | 0.73 |
|  |  | Quarterly | 14 | 0.60 | 6 | 9 | 0.58 | 10 | 0.58 | 4 | 8 | 0.65 |
|  |  | Annual | 14 | 0.58 | 6 | 9 | 0.58 | 9 | 0.54 | 4 | 9 | 0.53 |

**Table 2.**
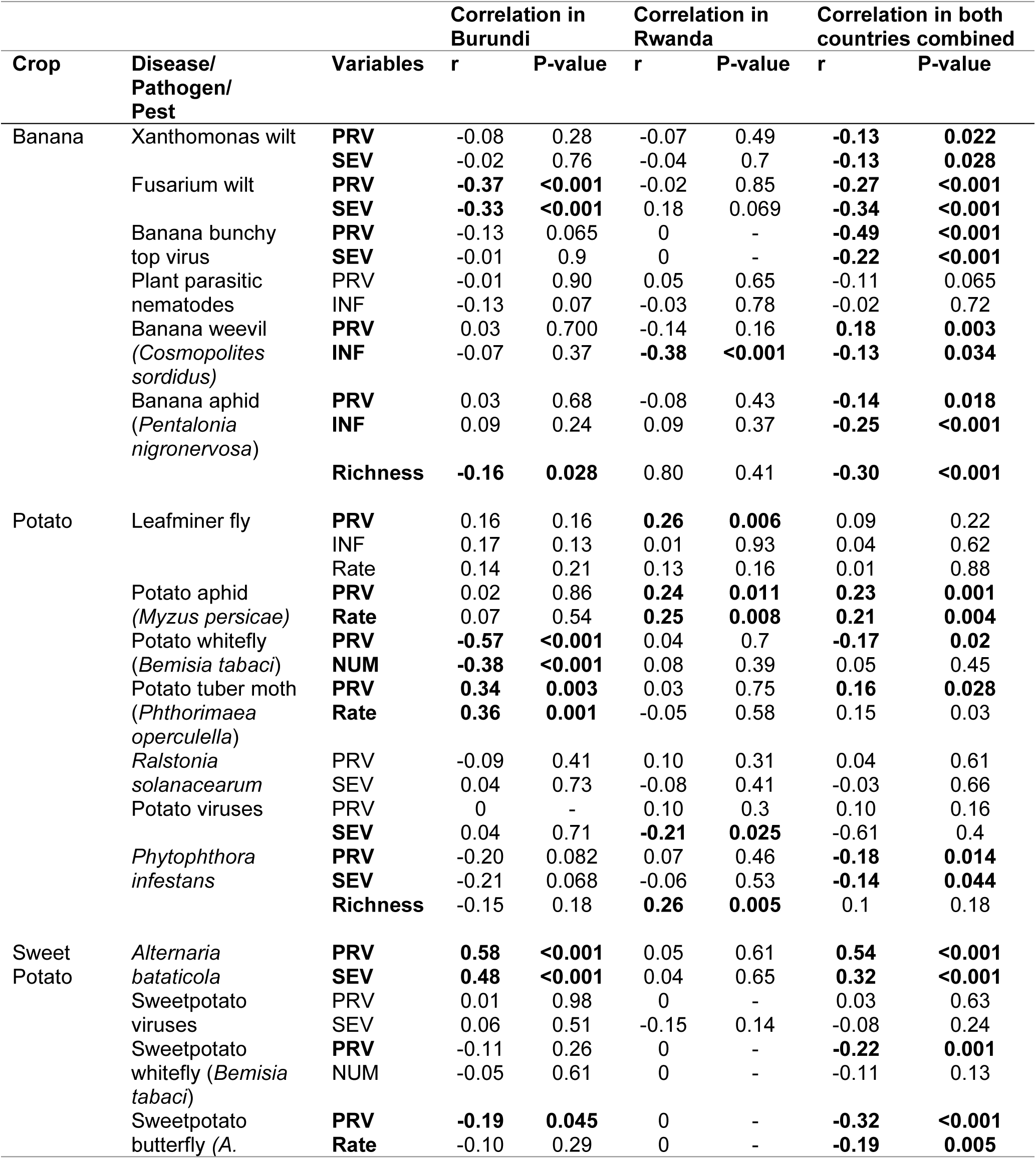
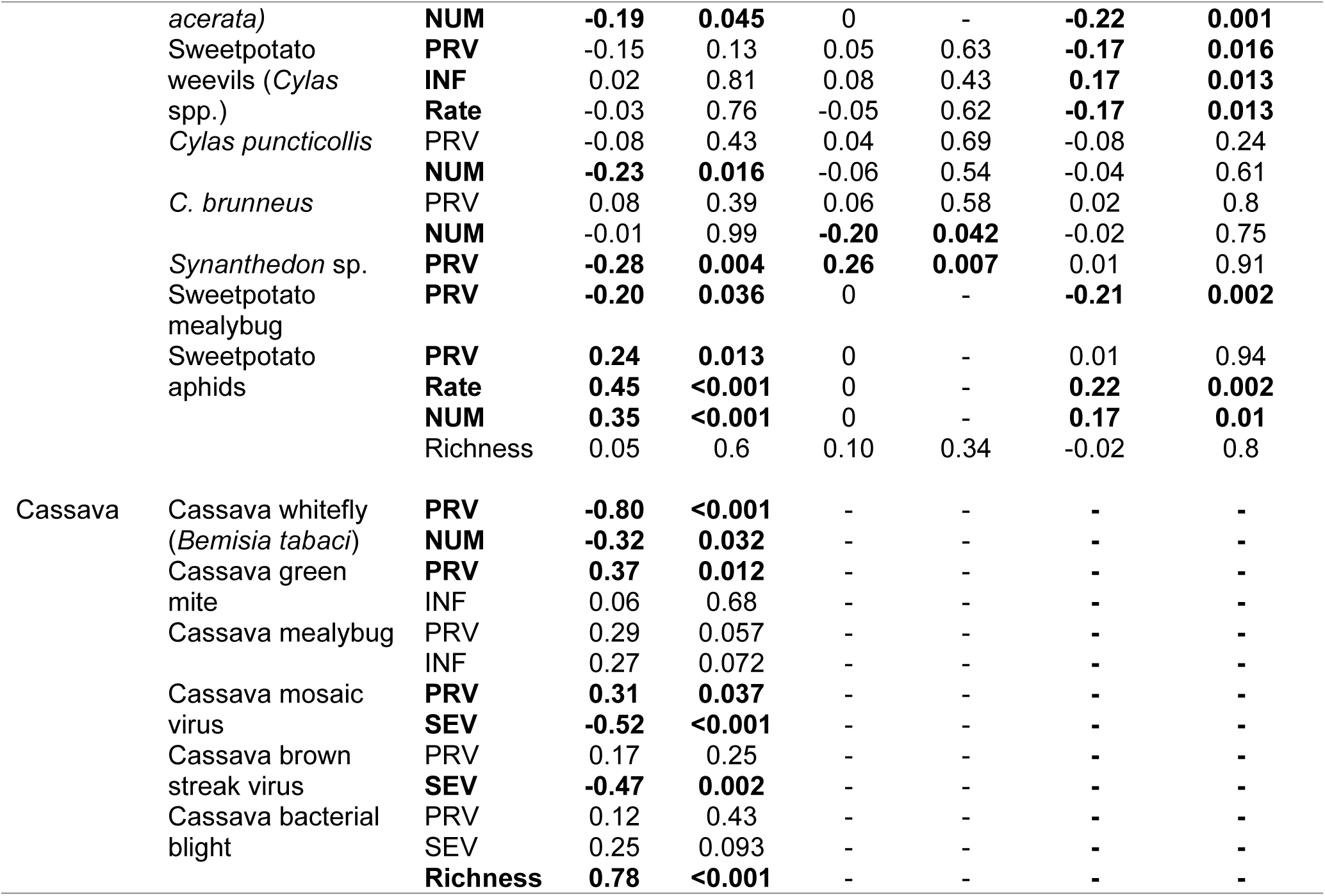
Correlation between altitude and pathogen and pest prevalence, severity, and infestation in Burundi and Rwanda (PRV: prevalence, SEV: severity, Rate: infestation rate, INF: infestation intensity, NUM: number of pests).

For banana, fields were surveyed in Burundi in March 2015 (during the rainy season) and in July 2016 (dry season), while fields were surveyed in Rwanda in July 2015 (dry season) and in November 2016 (rainy season). Cassava fields were surveyed only once in Burundi in March 2015 (rainy season), for reasons described above. For potato, field sampling was conducted in Burundi in July 2016 (dry season) and December 2017 (rainy season), and in Rwanda in June-July 2015 (dry season) and November-December 2017 (rainy season). For sweetpotato, the field surveys took place in Burundi in November-December 2015 (rainy season) and August 2016 (dry season), and in Rwanda in July 2015 (dry season) and November 2017 (rainy season). The irregular dates for these crop-field surveys were due in part to the civil unrest prevailing in the region at the time the study was conducted.

#### 2.2.2. Quantifying patterns of field-scale association of crop pathogens and pests

We evaluated the patterns of association of pathogens and pests based on prevalence and severity/infestation using non-metric multidimensional scaling (NMDS) (Kruskal, 1964) and heatmaps (Pryke *et al*., 2007) for visualization. We initially calculated five measures of dissimilarity (Bray, Manhattan, Jaccard, Chao, and robust-Aitchison) in the NMDS analysis using the ggvegan package v0.1.999 (Simpson, 2020) in R. We selected the Bray distance in the final analysis due to its highest goodness of fit. A heatmap was prepared using the metan package v1.18.0 (Olivoto and Lúcio, 2020) in R to map the association of pathogens and pests based on prevalence and severity/infestation rates.

### 2.3. Prediction of pathogen and pest damage and geographic range shifts

#### 2.3.1. Machine learning to predict pathogen and pest levels

Weather, climate, and geographic variables were used to predict severity or infestation rates/intensity, comparing machine learning algorithms. We retrieved weather variables (monthly, quarterly, and annual temperature, precipitation, and relative humidity) from the NASAPOWER database using the nasapower package v4.0.10 (Sparks, 2018) in R. The climate variables included the 19 bioclimatic variables, solar radiation (kJ m-2 day-1), wind speed (m s-1), and water vapor pressure (kPa) described, along with the altitude variable, in section 2.1.2. The CCRIs and cropland density were assembled as described in section 2.1.1.

Data were preprocessed to exclude collinearity between variables. A stratified split was then used to divide the data into training and test sets in an 8:2 ratio. We then trained ten machine-learning algorithms: logistic regression (GLM and GLMNET), support vector machine (SVM, with linear and radial basis function), neural network (NN), k-nearest neighbors (KNN), stochastic gradient boosting (GBM, generalized boosted modeling), bagged trees, classification and regression trees (CART), and random forest (RF). We compared the performance of these algorithms to predict the severity or infestation of each pathogen and pest using the caret package v6.0-94 (Kuhn, 2008) in R. We selected SVM (with radial basis function) and RF for further analyses based on their having the highest performance in terms of MAE (mean absolute error), RMSE (root mean squared error) and R-squared (Kaundal *et al*., 2006).

In the next stage using the SVM and RF models, we executed a 10-fold cross-validation with ten repetitions to prevent overfitting the training model and used a grid search for hyperparameter tuning. The tuning process was driven by trainControl, with RMSE-based model evaluation to prioritize accuracy. Hyperparameter tuning for SVM included C (regularization parameter) and Gamma (kernel coefficient), both with a tuneLength of 10. Hyperparameter tuning for RF included mtry (the number of predictors randomly selected at each division) with a tuneLength of 10. We trained the model using the train function in the caret package and estimated the performance parameters MAE, RMSE, and R-squared. We used the resulting model to predict severity or infestation rate/intensity for the test data and estimated the performance parameters (RMSE and R-squared).

#### 2.3.2. Pathogen and pest communities across altitudinal gradients

To study the geographic range of pathogen and pest communities associated with the four crops in the sample transects in Rwanda and Burundi, we evaluated how pathogen and pest levels (prevalence, disease severity or infestation rate/intensity) changed with altitude across the sampling locations. We also evaluated the pathogen and pest richness (number of pathogens and pests present from those included in the sampling) in communities during the dry and rainy seasons, across the altitude gradients. The associations between prevalence, infestation/severity and richness of pathogens and pests across altitude gradients were evaluated based on Pearson’s correlation.

#### 2.3.3. Climate change: impact on regional temperature patterns

We assessed potential climate shifts along the altitude gradients in the Great Lakes region, using current and future temperature data from WorldClim database v2.1 (Fick and Hijmans, 2017). We used future climatic variables from three global circulation models (GCMs: GISS-E2-1-G, MIROC6 and CNRM-CM6-1), two Shared Socioeconomic Pathways (SSP1-2.6 and SSP5-8.5), and four future time-periods: 2021-2040, 2041-2060, 2061-2080, and 2081-2100. We selected these GCM modeling groups because of their equilibrium climate sensitivity (ECS) (Hausfather, 2019). ECS indicates the severity of future warming impacts. CNRM-CM6-1 assumes a high value of ECS, 4.9 C, while GISS-E2-1-G and MIROC6 assumes low values of ECS, 2.7 and 2.6 respectively (Hausfather, 2019). The two SSPs used in this study represent different emission pathways by the year 2100: sustainability (SSP1-2.6) and fossil-fuel development (SSP5-8.5) emission scenarios featured by the radiation forcing of 2.6 and 8.5 W/m², respectively (Pinnegar *et al*., 2021). We calculated the average annual temperature across the three selected GCMs for each future time-period and SSP combination. We used the resulting density plots to assess potential shifts in the distribution of annual mean temperature at low (<1500 m.a.s.l.) and high (>1500 m.a.s.l.) altitudes, using the ggplot2 package v3.5.0 (Wickham, 2016) in R.

## 3. Results

### 3.1. Landscape structures influencing pathogen and pest risk

#### 3.1.1. Identifying locations with high cropland connectivity

The analysis of cropland connectivity in the Great Lakes region (i) quantifies how the geographic distribution of crop production could influence the potential spread of crop-specific pathogens and pests and (ii) identifies candidate priority locations for pathogen and pest surveillance for each food security crop (Fig. 2). These maps of cropland connectivity highlight locations in the region that are likely important for understanding and managing the risk of crop-specific pathogens and pests.

**Fig. 2.**
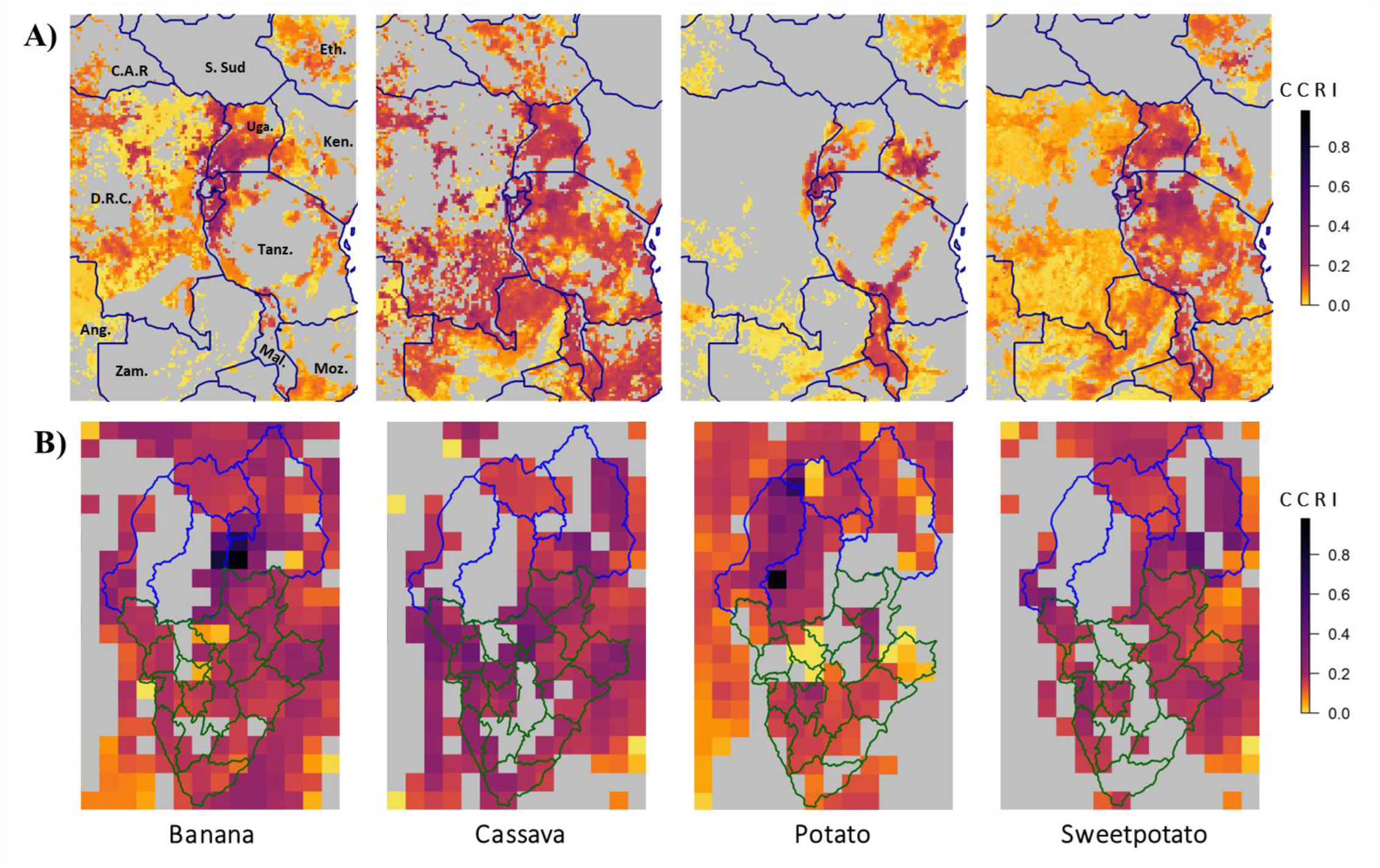
Maps of a cropland connectivity risk index (CCRI-betweenness) for four food security crops in (A) the Great Lakes region of Africa and (B) in a focused geographical extent in Rwanda and Burundi. The CCRI indicates how important a location is likely to be for potential spread of a host-specific pathogen or pest, based on the network of host availability. The CCRI is a weighted mean of four network metrics, including betweenness centrality in the results shown here. In (A), dark blue lines indicate national boundaries. In (B), blue lines indicate the provinces of Rwanda and green lines indicate the provinces of Burundi.

In these maps of cropland connectivity, a higher CCRI-betweenness (Fig. 2A) or CCRI-closeness indicates the potential to play an important role in the establishment and spread of crop-specific pathogens and pests affecting these crops. Locations with high CCRI are candidate priorities for surveillance and mitigation programs. Although CCRI-betweenness and CCRI-closeness may not yield identical CCRI values, they tended to highlight the same potential locations for prioritization (Fig. 2A). For banana, locations in Rwanda, Burundi, the DR Congo, Uganda, Tanzania, Ethiopia, Malawi, and Mozambique have a high CCRI (Fig. 2). Locations in the southern, eastern, and northern provinces of Rwanda have the highest CCRI-betweenness (Fig. 2A and 2B). For cassava, several locations across the region have a high CCRI. In Burundi, locations in the Cibitoke, Bubanza, Kirundo, Ngozi, Bujumbura Rural and Gitega provinces have the highest CCRI (Fig. 2A and B). For potato, locations in Rwanda, Burundi, the DR Congo, Uganda, Tanzania, Ethiopia, and Malawi have high CCRI-betweenness (Fig. 2). These maps also show that locations in the northern, western, and southern provinces of Rwanda have the highest CCRI-betweenness (Fig. 2A and B). For sweetpotato, locations in Rwanda, Burundi, Uganda, Tanzania, Malawi, and Ethiopia have a high CCRI (Fig. 2). These maps indicate the important role that Rwanda and Burundi play for understanding and mitigating the risks associated with the spread of pathogens and pests of these crops.

Maps of cropland connectivity (Fig. 2B) also illustrate the potential roles of the locations we selected for field sampling crop pathogens and pests in the Ruhengeri watershed in Rwanda and in the Rusizi watershed in Burundi. The connectivity of these landscapes varies from crop to crop. Some sampled locations had high cropland connectivity (e.g., cassava and sweetpotato in Burundi transects and potato in Rwanda transects) compared to other locations. Overall, the surveyed locations around Lake Kivu in Rwanda and Lake Tanganyika in Burundi, as well as other locations in the two countries, are likely to be important candidate locations for surveillance for the establishment and spread of banana, cassava, potato, and sweetpotato pathogens and pests in both countries, based on cropland connectivity (Fig. 2B).

#### 3.1.2. Associations among altitude, landscape structure and climate

The associations among (a) altitude, (b) crop landscape structure, and (c) climate variables such as temperature and precipitation, were analyzed for the Great Lakes region. Higher temperatures were generally associated with lower altitudes. There was a strong negative correlation between altitude and most temperature-related bioclimatic variables (Tem, Tmin, TColQ, TDriQ, TWetQ, TWarQ, Tmax, TColdQ). In contrast, altitude was weakly positively correlated with some precipitation-related bioclimatic variables such as PrecS, PWetQ, and PWetM, and negatively correlated with others, including Prec, PDriQ, PWarQ, and PColQ (Fig. 3). In this region, higher cropland density and cropland connectivity of banana, cassava, potato, and sweetpotato (Fig. 3) were at higher altitudes (except cassava density). Cropland connectivity of these crop species had a stronger positive correlation with altitude compared to cropland density. Cropland density and cropland connectivity for each of these four crops were negatively correlated with most temperature-related bioclimatic variables and positively correlated with most precipitation-related bioclimatic variables (Fig. 3).

**Fig. 3.**
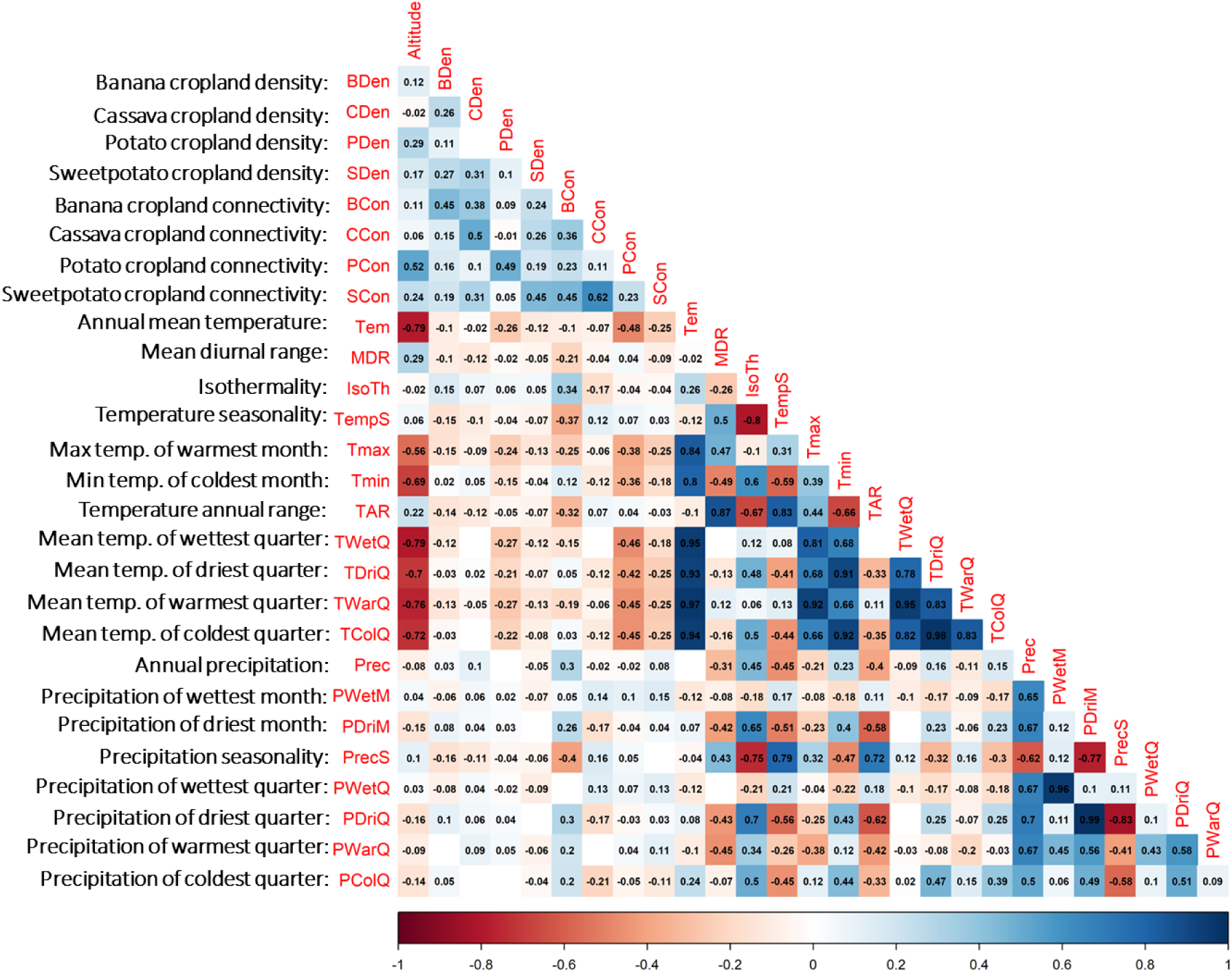
Associations between altitude, bioclimatic variables, and crop landscape structure (crop density and cropland connectivity) in the Great Lakes region of Africa, based on Pearson’s correlation. Strong positive correlations are in dark blue, while strong negative correlations are in dark red. White boxes without labels indicate no association.

### 3.2. Pathogen and pest communities

#### 3.2.1. Spatiotemporal structure of pathogen and pest communities

Most of the targeted pathogens and pests were found in Burundi and Rwanda in both crop-growing seasons (Fig. 4). Sweetpotato mealybugs, sweetpotato butterfly (*Acraea acerata*) and sweetpotato aphids were observed only in Burundi (Fig. 4). The observed prevalence of each pest or disease often varied by country, crop, and season. In both countries and seasons, the highest prevalence was observed for *Ralstonia solanacearum* and virus symptoms in potato, virus symptoms in sweetpotato, and *Cosmopolites sordidus* in banana.

**Fig. 4.**
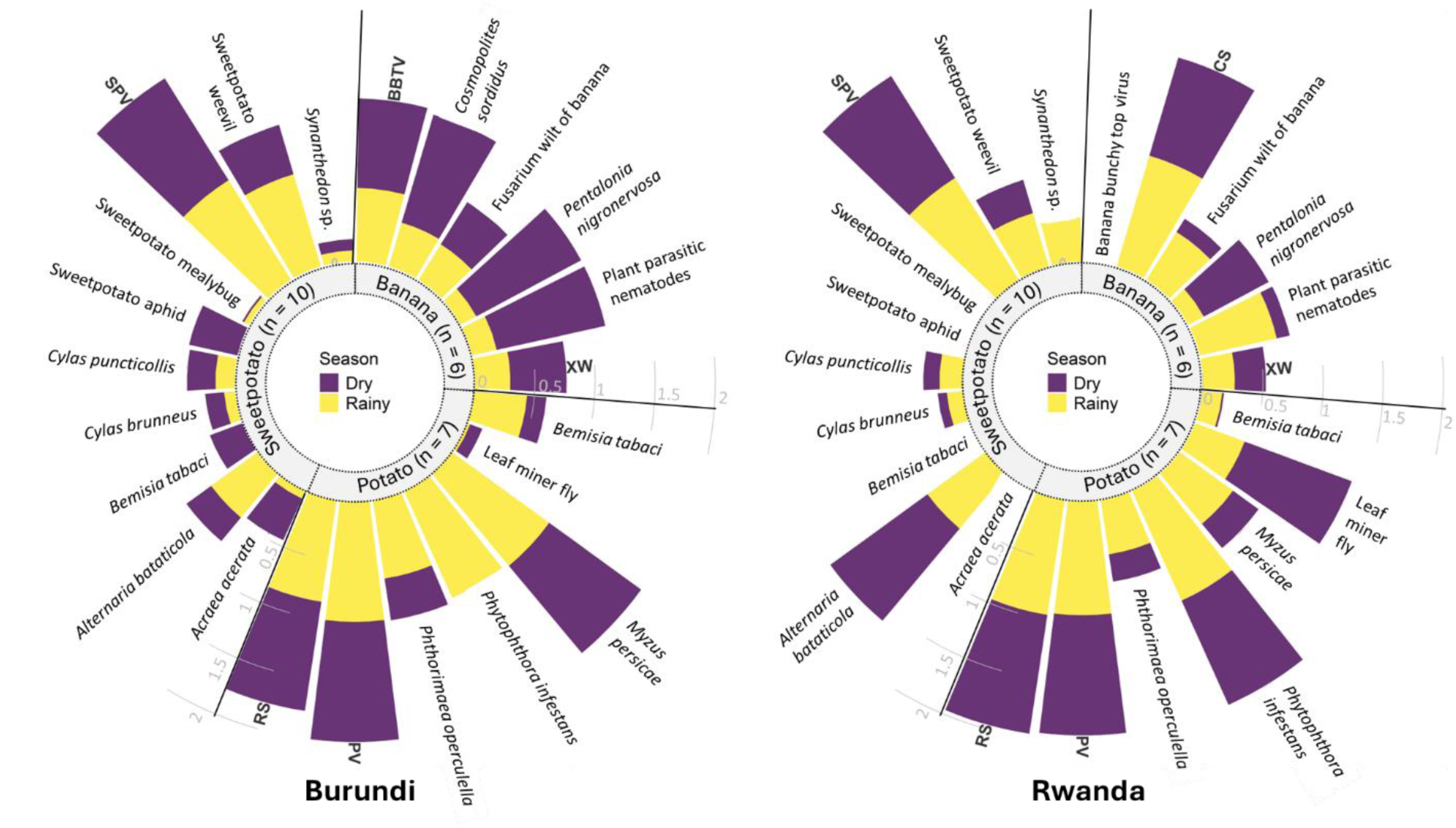
Seasonal distribution of observed pathogens and pests in smallholder production of banana, potato and sweetpotato in Burundi and Rwanda. (Cassava data are not represented here). The length of the bars represents the prevalence for both seasons (where a total of 2 indicates presence in every field in both seasons). Abbreviations: Banana bunchy top virus (BBTV), *Cosmopolites sordidus* (CS), *Ralstonia solanacearum* (RS), sweetpotato virus (SPV), and Xanthomonas wilt of banana (XW).

#### 3.2.2. Associations among pathogens and pests

The NMDS analysis identified a limited set of pathogens/pests whose levels were associated, although some were common enough that they frequently co-occurred with others (Fig. 5). A few pathogens and pests were associated in terms of prevalence and severity/infestation in the crop fields. Among banana diseases/pests, Fusarium wilt, Xanthomonas wilt and plant parasitic nematodes were associated in terms of both prevalence and severity (Fig. 5). In cassava, all pathogens and pests except *B. tabaci* were associated in terms of prevalence. However, *B. tabaci* was slightly associated with CMV in terms of infestation, which is expected as *B. tabaci* vectors CMV (Fig. 5). In potato, viruses and *R. solanacearum* were associated in terms of both prevalence and severity/infestation (Fig. 5). In sweetpotato fields, *B*. *tabaci* and sweetpotato aphids were associated, having similar infestation levels across fields.

**Fig. 5.**
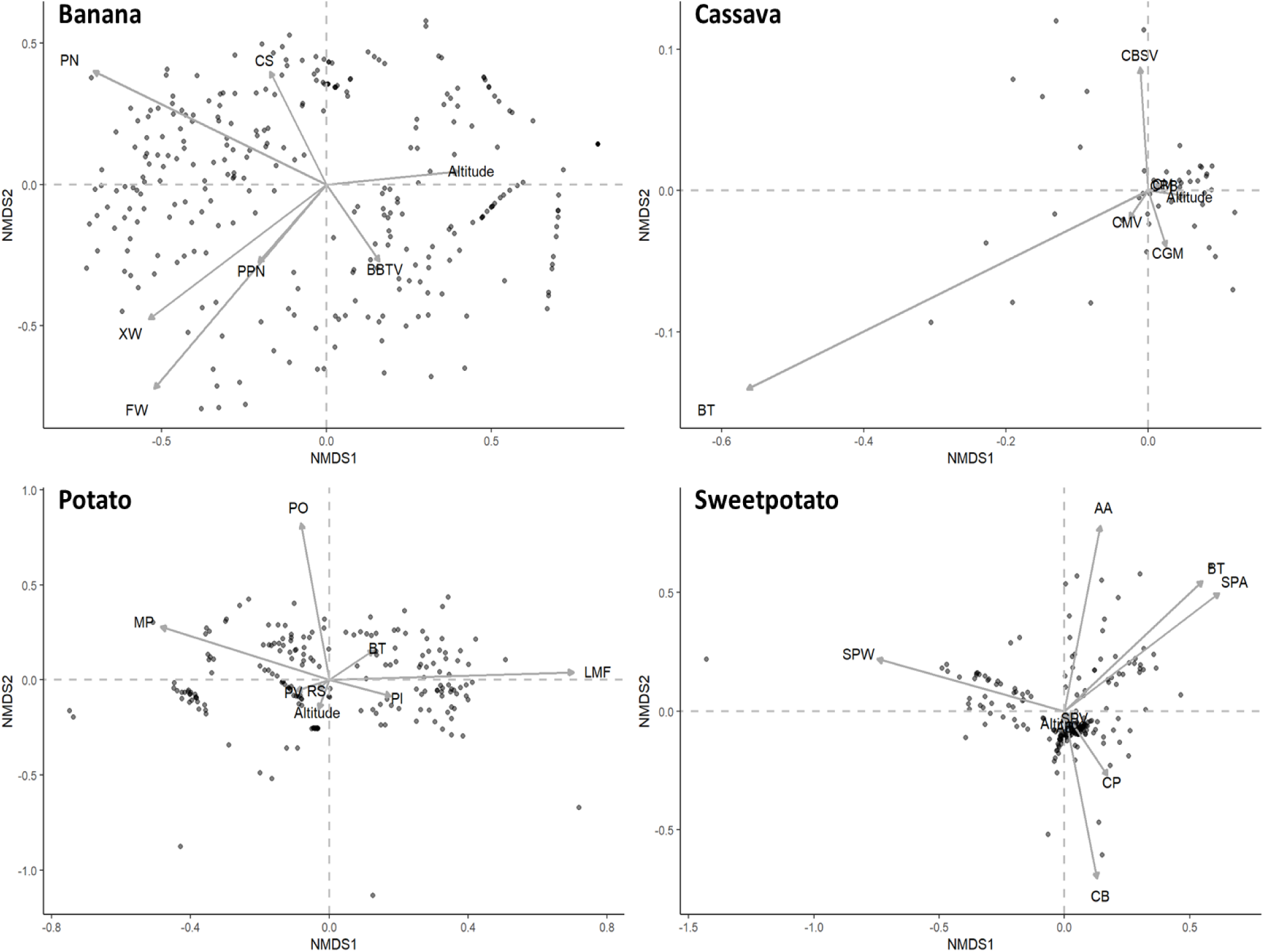
Relationships among economically important pathogens and pests and altitude in four major crops in Burundi and Rwanda, using non-metric multidimensional scaling (NMDS) based on Bray’s distance dissimilarities of disease severity or infestation intensity. Each dot represents a surveyed crop field. A line (or vector) indicates whether (line direction) and how strongly (line length) the pathogen or pest is associated with others. Nearly parallel lines in the same quadrant indicate pathogens and pests often co-occur in a crop. (Cassava data was collected only in Burundi for one season). Abbreviations: *Acraea acerata* (AA), *Alternaria bataticola* (AB), Banana bunchy top virus (BBTV), *Bemisia tabaci* (cassava, potato or sweetpotato whitefly; BT), cassava bacterial blight (CBB), Cassava brown streak virus (CBSV), cassava green mite (CGM), cassava mealybug (CM), Cassava mosaic virus (CMV), *Cylas brunneus* (CB), *Cylas puncticollis* (CP), *Cosmopolites sordidus* (CS), Fusarium wilt of banana (FW), leaf miner fly (LMF), *Myzus persicae* (MP), *Phytophthora infestans* (PI), plant parasitic nematodes (PPN), *Pentalonia nigronervosa* (PN), *Phthorimaea operculella* (PO), potato viruses (PV), *Ralstonia solanacearum* (RS), *Synanthedon* sp. (clearwing moth; Ssp), sweetpotato mealybug (SPM), sweetpotato virus (SPV), sweetpotato weevils (SPW), and Xanthomonas wilt of banana (XW).

Analyses of associations between pairs of pathogens/pests provide a more detailed perspective on which syndromes are common. In banana, some pairs like *C. sordidus* and *P. nigronervosa*, plant parasitic nematodes and Fusarium wilt, plant parasitic nematodes and *P. nigronervosa*, *P. nigronervosa* and Xanthomonas wilt, and Fusarium wilt and Xanthomonas wilt were positively correlated (p < 0.05) in terms of both prevalence and severity/infestation, while other pairs were only positively correlated (p < 0.05) in terms of prevalence: *C. sordidus* and plant parasitic nematodes, plant parasitic nematodes and Xanthomonas wilt, plant parasitic nematodes and BBTV, *P. nigronervosa* and BBTV, and BBTV and Xanthomonas wilt (Fig. 6).

**Fig. 6.**
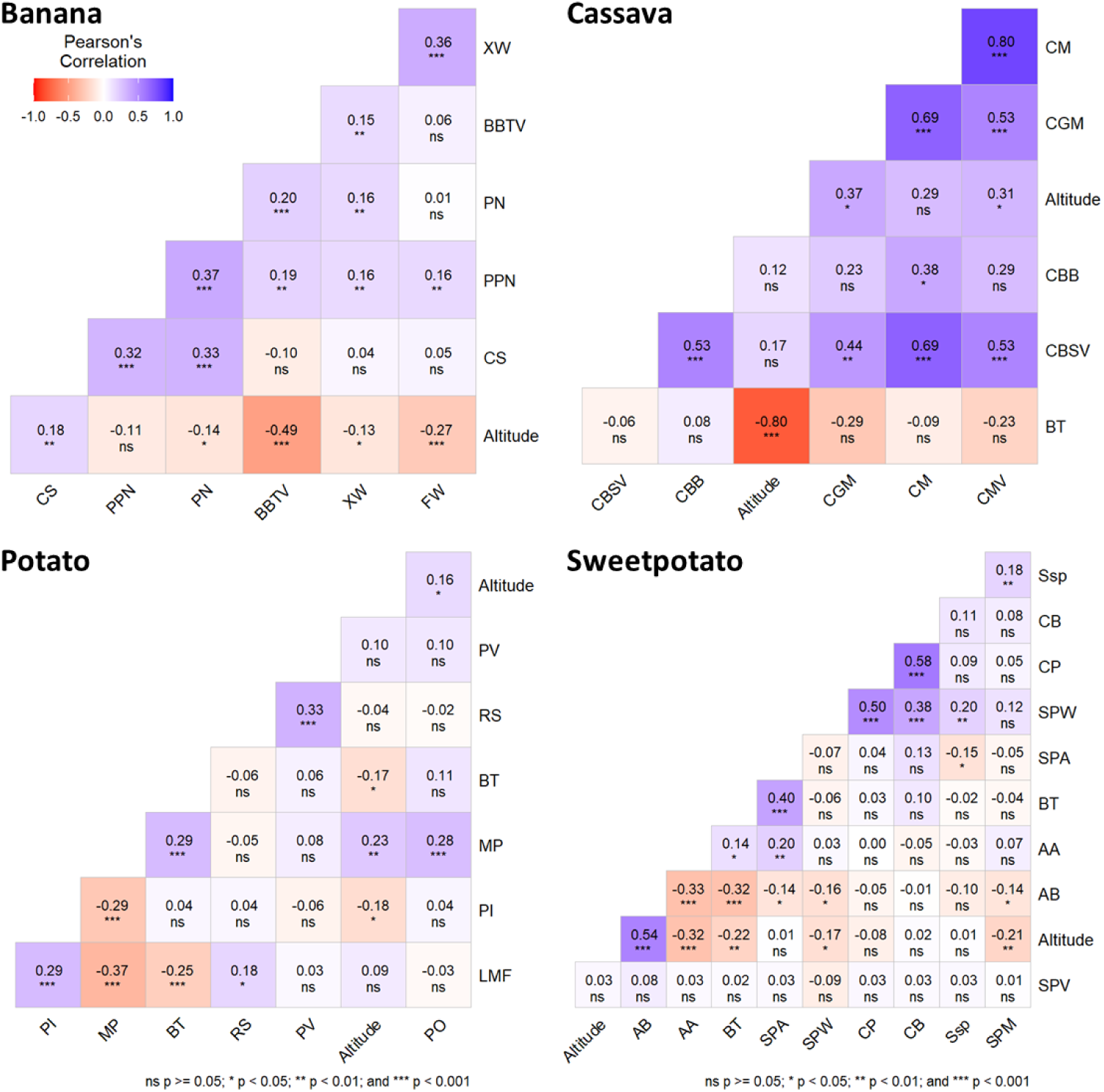
Association of pathogens and pests in banana, cassava, potato, and sweetpotato in smallholder farms in Burundi and Rwanda, based on Pearson’s correlation of prevalence. (Cassava data was collected only in Burundi for one season). Abbreviations: *Acraea acerata* (AA), *Alternaria bataticola* (AB), Banana bunchy top virus (BBTV), *Bemisia tabaci* (cassava, potato or sweetpotato whitefly; BT), cassava bacterial blight (CBB), Cassava brown streak virus (CBSV), cassava green mite (CGM), cassava mealybug (CM), Cassava mosaic virus (CMV), *Cylas brunneus* (CB), *Cylas puncticollis* (CP), *Cosmopolites sordidus* (CS), Fusarium wilt of banana (FW), leaf miner fly (LMF), *Myzus persicae* (MP), *Phytophthora infestans* (PI), plant parasitic nematodes (PPN), *Pentalonia nigronervosa* (PN), *Phthorimaea operculella* (PO), potato viruses (PV), *Ralstonia solanacearum* (RS), *Synanthedon* sp. (clearwing moth; Ssp), sweetpotato mealybug (SPM), sweetpotato virus (SPV), sweetpotato weevil (SPW), and Xanthomonas wilt of banana (XW).

In cassava, CBSV and mealybug were positively correlated (r = 0.69, p < 0.001 in terms of prevalence but negatively correlated (r = -0.44, p < 0.01) in terms of severity/infestation (Fig. 6). Only *B. tabaci* and CMV were positively correlated in terms of severity/infestation (r = 0.64, p < 0.001) while other pathogens and pests were positively correlated in terms of prevalence (p < 0.05). In potato, viruses and *R. solanacearum* was positively correlated in terms of prevalence (r = 0.33, p < 0.001) and severity (r = 0.21, p < 0.01), while *P. infestans* and *M. persicae*, and leaf miner fly and *M. persicae* were negatively correlated (p < 0.05) in terms of prevalence and severity/infestation, and leaf miner fly and *R. solanacearum* were positively correlated (p < 0.05) in terms of prevalence but negatively correlated (p < 0.05) in terms of severity/infestation (Fig. 6). In sweetpotato, *C. brunneus* and *C. puncticollis* were positively correlated in terms of prevalence (r = 0.58, p < 0.001) and infestation (r = 0.18, p < 0.01), while *A. bataticola* and weevils (*Cylas* spp.) were negatively correlated in terms of prevalence (r = -0.16, p < 0.05) but positively correlated in terms of severity/infestation (r = 0.18, p < 0.05) (Fig. 6). *A. bataticola* and mealybug, *A. bataticola* and aphids, *A. bataticola* and *B. tabaci*, *A. bataticola* and *Acraea acerata*, and aphids and *Synanthedon* sp. were negatively correlated in terms of prevalence (Fig. 6).

### 3.3. Prediction of pathogen and pest damage and geographic range shifts

#### 3.3.1. Machine learning to predict individual pathogen and pest levels

We compared machine learning algorithms in terms of their ability to predict severity and infestation in the field samples at monthly, quarterly, and annual intervals (Table 1). SVM and RF had the highest model performance for some pathogens/pests. For the SVM model, the performance characteristics across all pathogens and pests were 3-16% MAE for training, 7-20% and 4-26% RMSE, and 0.20-0.72 and 0.09-0.72 R-squared for training and testing, respectively, compared to the RF model with 3-17% MAE for training, 8-21% and 5-23% RMSE, and 0.23-0.72 and 0.12-0.73 R-squared for training and testing, respectively (Table 1). Among the 27 pathogens and pests tested, the three with the highest predictive performance metrics per crop were reported in Table 1.

The predictive performance was good for certain pathogens and pests, but poor for others (Table 1). As an example, SVM and RF performed well in predicting the monthly, quarterly, and annual infestation intensity of leaf miner fly in this region, explaining 69-71% of the variability in the data (R-squared) with low errors. The RMSE was 17-18% during training and 16-17% during prediction, demonstrating that both models generalized well to unseen data. In addition, the MAE of 11-13% indicated a low average error in training. The tight alignment between training and prediction metrics showed the robustness, stability and reliability of these models to predict leaf miner fly infestation intensity.

Pathogens and pests differed in the importance of predictor variables. Contemporary weather variables, such as monthly temperature, precipitation, and relative humidity (extracted through NASAPOWER), generally had high contribution scores. Cropland structure variables also played significant roles, with contribution scores reaching 100 for cropland density, 81 for cropland connectivity, and 69 for altitude. For example, the key variables for predicting leaf miner fly infestation intensity (monthly) using SVM were monthly temperature, precipitation, and relative humidity, precipitation in the driest month, cropland connectivity, cropland density and annual temperature range (i.e. difference between the maximum temperature of warmest month and the minimum temperature of coldest month).

#### 3.3.2. Pathogen and pest communities across altitudinal gradients

For each crop, pathogen and pest communities differed at low versus high altitudes (Table 2). Although there was not strong evidence for differences across the altitudinal gradients for 5 out of 27 pathogens and pests (correlations with p > 0.05 in Table 1), likely due in some cases to high variability across samples, there was evidence for differences in some pathogens and pests that distinguished the pathogen/pest communities (correlations in Table 2 with lower p-values). For instance, crop production at higher altitudes (>1,500 m.a.s.l) had more *Myzus persicae*, *Phthorimaea operculella*, and cassava green mites. At lower altitudes, crop fields had more Fusarium wilt and Xanthomonas wilt of bananas, *P. infestans*, *A. acerata* and *Bemisia tabaci* (Table 2). The distribution of these pathogens and pests across altitudes was supported in the analysis combining observations from Burundi and Rwanda; overall, for banana, potato, and sweetpotato, there was evidence that 44% of the targeted pathogens and pests were more common at low altitudes, and that 17% were more common at higher altitudes.

When analyzing data for Rwanda and Burundi separately, differences across altitudes for pathogens and pests were specific to the measure being used (e.g., disease intensity; Table 2) or the country. For example, potato fields at high altitudes had a higher prevalence of potato leaf miner fly, while there was not evidence for a difference in the infestation intensity of this pest across the altitudinal gradient. Similarly, there was stronger evidence that altitude was negatively correlated with the prevalence and severity of banana Fusarium wilt in Burundi (p < 0.001) compared to Rwanda (p = 0.069 for severity and p = 0.85 for prevalence). At the community level, potato fields in Rwanda and cassava fields in Burundi had higher pathogen and pest richness at higher altitudes, while richer pathogen and pest communities were found in banana fields at lower altitudes.

#### 3.3.3. Climate change and impact on regional temperature patterns

The frequency distribution of annual temperatures across altitudes in the Great Lakes region illustrates the transformative impact of climate change (Fig. 7). A progressive shift in temperature distribution is expected, highlighting how currently cooler climates at higher altitudes are likely to experience much higher temperatures (Fig. 7). A general warming trend is projected at both high and low altitudes. There is a greater temperature increase in the SSP5-8.5 scenario than in the SSP1-2.6 scenario. Currently, the mean annual temperature in the region ranges between 17-30°C at low altitudes, with the mode around 24°C, and between 10-22°C at higher altitudes, with the mode around 17°C. As climate change progresses, and depending on socio-economic pathways, this temperature landscape is likely to change substantially. Under the SSP1-2.6 scenario, by 2090 (2081-2100), temperatures are expected to range between 20-32°C at low altitudes, with the mode around 26°C, and between 10-25°C at high altitudes, with mode 20°C. The high-emission scenario (SSP5-8.5) predicts that temperatures will range between 22-36°C at low altitudes, with the mode around 28°C, and between 12-30°C at higher altitudes, with the mode around 25°C.

**Fig. 7.**
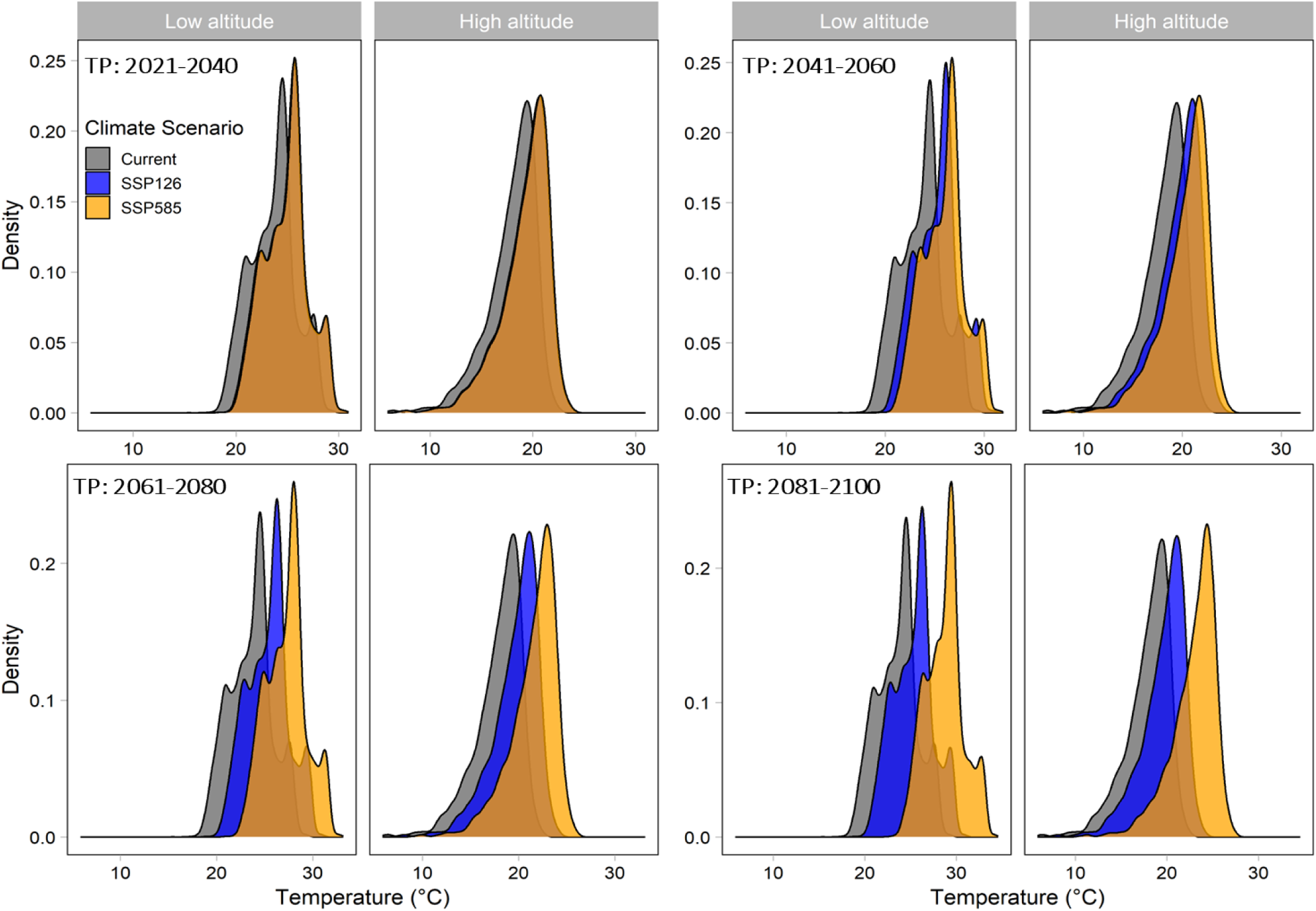
Shift in annual temperature range at lower and higher elevations in the Great Lake region of Africa. Density is the probability density or likelihood of temperature rates occurring at locations in the Great Lakes region of Africa. TP: time period of the projections (while the ‘Current’ reference remains the same). SSP: Shared Socioeconomic Pathway, sustainability (SSP1-2.6) and fossil-fueled development (SSP5-8.5) emission scenarios. Here, high altitude refers to locations above 1500 m.a.s.l. and low altitude those below 1500 m.a.s.l.

These projections suggest future climate conditions at higher altitudes could become more favorable for the 44% of pathogens and pests we evaluated that currently are more common at lower altitudes (section 3.3.2). Likewise, pathogens and pests currently more common at higher altitudes (17% of those evaluated in section 3.3.2) may become less common as the locations warm in the future. These percentages are based on banana, potato, and sweetpotato, the three crops for which we had the most complete data sets for analysis in predictive models based on field data collected to allow direct comparisons across pathogens and pests.

## 4. Discussion

This study identified environmental predictors for a set of important pathogens and pests in tropical agroecosystems key to food security. We identified candidate surveillance locations based on cropland (host) connectivity, a starting point for understanding geographic spread and establishment of host-specific pathogens and pests. Based on the current geographic and seasonal distribution of 27 economically important pathogens and pests across climate gradients, we evaluated future climate scenarios. The 44% of the pathogens and pests currently more abundant at lower altitudes could become more common at higher altitudes under climate change, while the 17% more abundant at higher altitudes could become less common overall in the future. Identifying differences in pathogen and pest communities and their predictors contributes to a more complete understanding of agroecosystems, which can be translated to inform management strategies, as we discuss below.

### 4.1. Pathogen and pest risk across landscape structures

The cropland connectivity analysis provides a first approximation to the geographic distribution of pathogen and pest risk, based on host availability. This analysis identified candidate priority locations for surveillance and mitigation of the potential spread of host-specific pathogens and pests based on networks of host availability (Fig. 2). Analysis of cropland connectivity provides a baseline for quantifying invasion risks for pathogens and pests at a national or global scale (Andersen Onofre *et al*., 2021; Mouafo-Tchinda *et al*., 2024; Etherton *et al*., 2025). In the Great Lakes region, locations in Rwanda and Burundi had a high CCRI for all four food-security crops, highlighting their vulnerability to pathogen and pest spread in the absence of phytosanitary measures. Maps of cropland connectivity can inform monitoring of outbreaks of introduced or emerging diseases and pests by national plant protection organizations (NPPOs). Ongoing improvements to understanding other components of pathogen and pest risk – such as the effects of climate change, local trade, genetic resistance deployment, and other management adoption – could be integrated in more complete habitat connectivity analyses as additional resources become available (Keshav *et al*., 2024).

### 4.2. Pathogen and pest communities

Analysis of the on-farm distribution of 27 pathogens and pests highlights how these communities change along climate gradients, in terms of seasonal abundance (Fig. 4), community structure (Fig. 5), and interspecific pathogen/pest associations (Fig. 6). Previous studies have typically addressed how individual tropical pathogens or pests are distributed along climate gradients (Ebregt *et al*., 2005; Okonya and Kroschel, 2013; Nyang’au *et al*., 2021); a community perspective provides a more realistic scenario for understanding regional pathogen and pest management (Savary *et al*., 2000; Were *et al*., 2013; Makiola *et al*., 2021). The study addresses this knowledge gap by identifying which pathogens/pests are most prevalent across seasons, which pathogen/pest occurrences are associated, and how climate gradients are likely to influence disease severity and pest infestation. Evaluating the interactions and cumulative effects of multiple pathogens/pests in specific production environments can increase our understanding of injury profiles (Savary *et al*., 2000), the cumulative damage to plants caused by the set of biotic stressors including insects, pathogens and weeds.

Our findings highlight injury profiles for food security crops across climate gradients. Some associations are already well-known; as expected, BBTV and its vector *P. nigronervosa* were positively correlated (Hu *et al*., 1996; Blomme *et al*., 2020). BBTV has been present in Africa for decades, its spread continues to have an impact on new regions, and its management remains a major challenge (Rybicki, 2015; Ngatat *et al*., 2024; Ocimati *et al*., 2024). We also found strong associations between certain pathogens and pests; Fusarium wilt, Xanthomonas wilt and plant parasitic nematodes were associated in banana, viruses and *R. solanacearum* in potato, and *B. tabaci* and aphids in sweetpotato. These pathogen/pest associations underscore the need for integrated management strategies that address the spread and combined impacts of multiple pathogens and pests on plants (Mouafo-Tchinda *et al*., 2022; Alcalá Briseño *et al*., 2023).

### 4.3. Climate-based predictions of pathogen and pest levels

In the Great Lakes region, temperature generally decreased with increasing altitude (Fig. 3), where higher temperatures were associated with an increase in levels of some pathogens and pests and a decrease in others (Table 2). Whiteflies were more common at low altitudes for cassava, potato and sweetpotato. Aphids were more common at low altitudes for banana, but more common at high altitudes for potato. The observed differences in the altitudinal distribution of these pests could be attributed to their species composition, host specificity, and the influence of species-specific ecological adaptations (Aleuy and Kutz, 2020; McCulloch and Waters, 2023). Aphid communities included different species, with *P. nigronervosa* more frequent in low-altitude plants, and *M. persicae* more frequent in high-altitude plants (Malumphy *et al*., 2017; Wells and Clark, 2019; Wieczorek *et al*., 2019; Aleuy and Kutz, 2020).

Weather variables are well known to be useful for predicting disease severity (Kaundal *et al*., 2006; Carisse and Fall, 2021; Fenu and Malloci, 2021). Cropland geography variables (cropland density, connectivity), altitude, and climate variables (annual mean temperature, temperature annual range, and annual precipitation) were key for most of the pathogens and pests we evaluated. Model performance depends on data availability, the predictor variables, and the biological features of pathogens and pests (Xiao *et al*., 2018; Fenu and Malloci, 2021). The data from Rwanda and Burundi supported prediction of some pathogens, while others, like BBTV, would require more study. The models effectively predicted leaf miner fly infestation intensity, accounting for 69-71% of the variability with low training and prediction errors. These models could serve as a prototype for predictive models of leaf miner fly infestation intensity in this region (Table 1).

Temperature influences the life cycle, reproduction, and migration patterns of pathogens and pests (Awmack *et al*., 1997; Yamamura and Kiritani, 1998; Mouafo-Tchinda *et al*., 2021). In response to regional warming, certain pathogens and pests may become more common at specific altitudes (Poveda *et al*., 2012; Datta *et al*., 2017; Blomme *et al*., 2020), which will require adaptation of pathogen and pest management to future conditions. Several studies suggest that climate change shifts some species ranges to higher altitudes (Bale *et al*., 2002; Colwell *et al*., 2008; Bebber *et al*., 2013), and the new results from this study clarify the specific effects on individual pathogens and pests in Rwanda and Burundi for key food security crops.

Pathogen/pest richness in banana decreased with altitude, while it increased in cassava. In addition, the severity of certain pathogens/diseases such as BBTV, CMV, *P. infestans* and Fusarium wilt decreased with altitude, while the severity or infestation of *A. bataticola* and aphids in sweetpotato increased. In general, we found evidence that 44% of the pathogens and pests currently more common at low altitude in banana, potato, and sweetpotato could shift to higher altitudes under climate change, based on climate matching. Specifically, those expected to become more common at higher altitudes are Xanthomonas wilt, Fusarium wilt, BBTV, and *P. nigronervosa*, in banana; sweetpotato mealybug, *A. acerata*, sweetpotato weevils in sweetpotato; and *B. tabaci* in potato and sweetpotato. The 17% of pathogens and pests currently more common at high altitudes could become less common. Cropland connectivity was positively correlated with altitude and was a key predictor of severity/infestation of some pathogens and pests. This suggests that food-security crops at higher altitudes may experience substantial flows of pathogens and pests, amplifying the risk of damage. Customizing risk assessments for these specific pest/pathogen communities across altitudes could make management strategies better adapted to the challenges posed by climate change for each crop in the region.

### 4.4. Translating agroecosystem understanding of pathogens and pests to support current and future agriculture

Historically, global and national research and extension efforts have been employed to manage diseases and pests in the Great Lakes region (RTB, 2013; Kroschel *et al*., 2014). Integrated management systems combining cultural control, surveillance, awareness creation and involving policy makers and farmers have been adopted (Legg *et al*., 2006; Gotor *et al*., 2022). There has been support for active management programs where clean planting material is provided to farmers to encourage them to destroy diseased plants. Efforts have been made to create awareness and empower farmers to recognize and destroy diseased plants. A challenge for these efforts is limited resources to support ongoing, sustainable plans by governments and national programs, especially after project lifespans are over. The results of this new study can help to inform prioritization and efficient use of available resources. Currently, for many food security crops, efforts are underway to improve seed health, obtain resistance to key pathogens and pests, and adapt these improved technologies and techniques for use in humanitarian settings (RTB, 2013; Andrade-Piedra *et al*., 2020; Andrade-Piedra *et al*., 2023).

Pathogen and pest risk analysis can help to ensure the economic, social, and environmental sustainability of crop production (Kaundal *et al*., 2006; Fall *et al*., 2016; Carisse and Fall, 2021). Image analysis apps such as the PlantVillage Nuru app and the Tumaini app can help smallholder farmers in the region diagnose plant damage caused by pests and diseases, and offer national plant protection organizations and extension staff a tool for surveillance and disease mapping (Ramcharan *et al*., 2019; Selvaraj *et al*., 2019; Kreuze *et al*., 2022). Predicting the timing of pathogen and pest threats would enable farmers to adjust management strategies in the short run and would support prioritization in research and breeding programs in the long run.

Ongoing improvements in predicting pathogen and pest impacts depend on weather data availability, which is often limited in low-income and disaster-prone countries. Satellite platforms like NASA POWER offer a useful alternative (Sparks, 2018), providing near-real-time weather data and allowing researchers in resource-limited settings to develop useful models without the need for extensive ground observations. Another challenge is detection of transient pests such as aphids, which move frequently and rapidly across fields (Loxdale, 2018). Innovative sampling techniques, such as suction traps and environmental DNA (eDNA) analysis, could be helpful for monitoring pest populations more accurately (Taberlet *et al*., 2012; Poppinga *et al*., 2016). However, implementing these advanced approaches in low-income and disaster-prone areas such as the Great Lakes region remains a challenge due to limited resources and technical capacity, requiring scalable and cost-effective solutions.

In disaster-prone regions, humanitarian organizations and government agencies must prioritize key areas where abandonment or poor management of staple crops is particularly important to regional spread of pathogens and pests (Etherton *et al*., 2024; Mouafo-Tchinda *et al*., 2024). This study identifies candidate priority locations for mitigation of the establishment and spread of pathogens and pests, characterizes pathogen and pest associations (injury profiles), and evaluates potential altitudinal shifts under the influence of climate change. Future research can build on these results by validating and refining the models for each pathogen/pest, and developing tools that growers and policymakers can easily implement. Ongoing improvement can increase the adaptability and effectiveness of plant health management strategies, a no-regrets adaptation strategy as agricultural systems face increasing challenges due to climate change.

## Acknowledgements

We appreciate support from the CGIAR Seed Equal Research Initiative, the CGIAR Roots Tubers and Bananas Research Program, USAID Bureau of Humanitarian Assistance (BHA) award number 720BHA22IO00136, the OneCGIAR Initiative on Plant Health, and the CGIAR Trust Fund (www.cgiar.org/funders/); we thank all donors and organizations which globally support the work of CGIAR through their contributions to the CGIAR Trust Fund. We also appreciate support from USDA Animal and Plant Health Inspection Service (APHIS) Cooperative Agreements AP21PPQS&T00C195 and AP22PPQS&T00C133. The opinions expressed in this article are those of the authors and do not necessarily reflect the view of USAID BHA or USDA APHIS.

